# Predictors of cognitive impairment in primary age-related tauopathy: an autopsy study

**DOI:** 10.1101/2021.06.08.447553

**Authors:** Megan A. Iida, Kurt Farrell, Jamie M. Walker, Timothy E. Richardson, Gabe Marx, Clare H. Bryce, Dushyant Purohit, Gai Ayalon, Thomas G. Beach, Eileen H. Bigio, Etty Cortes, Marla Gearing, Vahram Haroutunian, Corey T. McMillan, Eddie B. Lee, Dennis Dickson, Ann C. McKee, Thor D. Stein, John Q. Trojanowski, Randall L. Woltjer, Gabor G. Kovacs, Julia K. Kofler, Jeffrey Kaye, Charles L. White, John F. Crary

**Affiliations:** Department of Pathology, Neuropathology Brain Bank & Research CoRE, Nash Family Department of Neuroscience, Friedman Brain Institute, Ronald M. Loeb Center for Alzheimer’s Disease, Icahn School of Medicine at Mount Sinai, USA; Department of Pathology and Laboratory Medicine and The Glenn Biggs Institute for Alzheimer’s & Neurodegenerative Diseases, UT Health San Antonio, San Antonio, TX, USA; Ultragenyx Pharmaceuticals USA; Neuropathology, Banner Sun Health Research Institute, Sun City, Arizona, USA; Department of Pathology, Northwestern Cognitive Neurology and Alzheimer Disease Center, Northwestern University Feinberg School of Medicine, Chicago, Illinois, USA; Department of Pathology and Laboratory Medicine, Emory University School of Medicine, Atlanta, Georgia, USA; Departments of Psychiatry and Neuroscience; Alzheimer’s Disease Research Center, Icahn School of Medicine at Mount Sinai, New York, New York, USA; and JJ Peters VA Medical Center (MIRECC), Bronx, NY; Department of Neurology, Perelman School of Medicine, Penn FTD Center, Center for Neurodegenerative Disease Research, University of Pennsylvania, Philadelphia, Pennsylvania, USA; Department of Neuroscience, Mayo Clinic, Jacksonville, Florida, USA; Department of Pathology, VA Medical Center & Boston University School of Medicine, Boston, Massachusetts, USA; Center for Neurodegenerative Disease Research, Department of Pathology and Laboratory Medicine, Perelman School of Medicine, University of Pennsylvania, Philadelphia, Pennsylvania, USA; Department of Pathology, Oregon Health Sciences University, Portland, Oregon, USA; Laboratory Medicine Program & Krembil Brain Institute University Health Network Toronto Ontario Canada; Tanz Centre for Research in Neurodegenerative Disease and Department of Laboratory Medicine and Pathobiology, University of Toronto, Toronto, Ontario, Canada; Previous address: Institute of Neurology, Medical University of Vienna, Vienna, Austria; Department of Pathology, University of Pittsburgh Medical Center, Pittsburgh, Pennsylvania, USA; Department of Neurology, Oregon Health & Science University, Portland USA; Neuropathology Laboratory, Department of Pathology, University of Texas Southwestern Medical Center, USA

**Keywords:** PART, dementia, Aging, Braak, ARTAG

## Abstract

Primary age-related tauopathy (PART) is a form of Alzheimer-type neurofibrillary degeneration occurring in the absence of amyloid-beta (Aβ) plaques. While PART shares some features with Alzheimer disease (AD), such as progressive accumulation of neurofibrillary tangle pathology in the medial temporal lobe and other brain regions, it does not progress extensively to neocortical regions. Given this restricted pathoanatomical pattern and variable symptomatology, there is a need to reexamine and improve upon how PART is neuropathologically assessed and staged. We performed a retrospective autopsy study in a collection (*n*=174) of post-mortem PART brains and used logistic regression to determine the extent to which a set of clinical and neuropathological features predict cognitive impairment. We compared Braak staging, which focuses on hierarchical neuroanatomical progression of AD tau and Aβ pathology, with quantitative assessments of neurofibrillary burden using computer-derived positive pixel counts on digitized whole slide images of sections stained immunohistochemically with antibodies targeting abnormal hyperphosphorylated tau (p-tau) in the entorhinal region and hippocampus. We also assessed other factors affecting cognition, including aging-related tau astrogliopathy (ARTAG) and atrophy. We found no association between Braak stage and cognitive impairment when controlling for age (*p*=0.76). In contrast, p-tau burden was significantly correlated with cognitive impairment even when adjusting for age (*p*=0.03). The strongest correlate of cognitive impairment was cerebrovascular disease, a well-known risk factor (*p*<0.0001), but other features including ARTAG (*p*=0.03) and hippocampal atrophy (*p*=0.04) were also associated. In contrast, sex, *APOE*, psychiatric illness, education, argyrophilic grains, and incidental Lewy bodies were not. These findings support the hypothesis that comorbid pathologies contribute to cognitive impairment in subjects with PART. Quantitative approaches beyond Braak staging are critical for advancing our understanding of the extent to which age-related tauopathy changes impact cognitive function.

## Introduction

It is widely recognized that abnormal hyperphosphorylated tau (p-tau) deposition is a ubiquitous feature of the aging human brain, observed in both cognitively normal subjects and in those with a range of clinical features, including cognitive, motor and psychiatric symptoms [37]. The causes of tauopathy are diverse, and include both genetic and environmental risk factors [48]. Autosomal dominant mutations in the tau gene (*MAPT*) cause frontotemporal lobar degeneration and common risk alleles, notably the *MAPT* 17q21.31 H1 haplotype, are associated with sporadic tauopathies including progressive supranuclear palsy (PSP), corticobasal degeneration (CBD), and argyrophilic grain disease (AGD) [12]. Abnormal p-tau deposition is also seen following exposure to repetitive head trauma in contact sports and other contexts in the setting of chronic traumatic encephalopathy (CTE) [43]. Neurofibrillary tangles (NFT) are also a component of Alzheimer disease (AD), where they are associated amyloid-beta deposits [16].

Although it is generally understood that autopsy studies are critical for establishing definitive diagnoses, the neuropathology of the tauopathies is complex and overlapping. Further, non-impaired individuals often display significant amounts of p-tau accumulation, complicating our understanding of the contribution of such brain changes to symptomatology. Approaches to assessing tauopathy in post-mortem tissues continue to evolve. Neuropathologically, tauopathies can be differentiated by the neuroanatomical regionality of p-tau aggregates, cell type involvement (i.e., neurons versus glia), preferential isoform accumulation, and filament ultrastructure. Based upon these differentiating features, validated neuropathological diagnostic consensus criteria have been devised and, in some cases, undergone revision. Examples include revision of the AD diagnostic criteria, and consensus criteria for CTE [41, 46].The term aging-related tau astrogliopathy (ARTAG), which was described in recent consensus criteria on various patterns of astrocytic p-tau observed in aging, has been especially helpful for differentiating age-related changes from CTE, both of which have perivascular p-tau deposits, but with differences in cell types involved [38, 42]. The introduction of criteria for primary age-related tauopathy (PART) to describe individuals who develop AD-type neurofibrillary pathology with or without dementia in the absence of significant amyloid deposition helped to better define this entity and differentiated it from AD [17]. Understanding age-related tauopathy is of critical importance in the context of diagnosis and staging of all the tauopathies given its extremely high prevalence and importance as a co-morbidity in essentially all studies evaluating tauopathy.

There has been controversy surrounding the PART consensus criteria since their introduction [11, 19], and there have been a substantial number of recent clinicopathological studies focused on understanding this pathological presentation [4, 6, 7, 29, 33, 36, 51, 52, 60]. Given the close clinical and neuropathological similarities between PART and AD such that historically the two entities were classified together, accumulating evidence has highlighted differences. Clinically, the average age is higher for individuals who have PART than those with AD and patients with PART are more often female [35]. Patients with PART pathology are more often cognitively normal, but a subset have mild cognitive impairment or amnestic dementia, and this correlates with p-tau severity [17]. Among symptomatic individuals with a neuropathological diagnosis of PART, nearly half had been clinically diagnosed with AD compared with 86% of those with autopsy-confirmed AD, indicating that despite diagnostic uncertainty, clinicians recognize differences between the two [59]. One retrospective study identified other factors including depression, Braak stage, and history of stroke, as independent predictors of cognitive impairment [6]. Another found that those with PART had a sparing of semantic memory compared to those with AD, suggesting that there is a distinct difference in clinical presentation [8]. Longitudinal analyses found that subjects with PART have a significantly slower clinical decline after becoming symptomatic than those with AD across multiple neuropsychological domains [60].

One limitation of most published studies on PART is that they rely on retrospective analysis of previously collected datasets (e.g., the National Alzheimer’s Coordinating Center database, NACC) with predefined neuropathological measures that may not fully capture all the clinically relevant features [45]. Further, findings might not be generalizable to other populations, and a lack of uniform analysis and quantitation might lead to bias. Critically, the Braak staging system was specifically developed for assessment of tau pathology in the context of AD, and has not been rigorously tested in amyloid-negative subjects, so the extent to which it is valid for staging p-tau pathology in PART is unclear. Additionally, the Braak stage represents a hierarchical progression of the regional spread of neurofibrillary tangles, but does not directly measure the severity or burden of p-tau, but this has been incorporated into some operationalized frameworks [2]. Because the pathology in PART generally remains predominantly in the medial temporal lobe, this hierarchical pathoanatomical system may sub-optimally measure severity of the disease. There are numerous approaches to assessing lesion burden of p-tau and other pathologies [10, 28, 30, 31, 40, 41, 44, 63], including cell counting and stereology [3, 5, 13, 21, 27, 64]. While each of these approaches have intrinsic advantages, they are limited in that they are labor intensive and for this reason and others, these methods have not been widely adopted in neuropathology laboratories [20, 62]. One approach that may have potential to better assess p-tau in PART is using computer-assisted quantitative morphometrics on digital whole slide images, which may be well suited for staging PART.

Here, we studied a cohort of autopsy-confirmed subjects with PART, enabling us to reexamine how tau pathology manifests in PART. We compared Braak staging with computer-assisted quantitative measures of p-tau burden, and used logistic regression to assess their contribution to cognitive impairment. Using this cohort, we were able to explore critical co-morbid pathologies (e.g., cerebrovascular disease), and further assess neuropathological changes that are not available in existing publicly available datasets, including atrophy and ARTAG.

## Methods

### Patient samples

Formalin-fixed paraffin embedded (FFPE) tissue from the frontal cortex and hippocampus as well as fresh-frozen tissue from frontal cortex were derived from autopsy brains from a subset of individuals from a previously described collection [61]. Specifically, the cohort included cases from the Oregon Health Sciences University (Portland, OR, USA), Banner Sun Health Research Institute (Sun City, AZ, USA), Emory (Atlanta, GA, USA), Northwestern (Evanston, IL, USA), the University of Pennsylvania (Philadelphia, PA, USA), University of Pittsburgh (Pittsburgh, PA, USA), University of Texas Southwestern Medical Center (Dallas, TX, USA), and the Medical University of Vienna (Vienna, Austria). Clinical inclusion criteria included being cognitively normal or having a diagnosis of mild cognitive impairment (MCI) or dementia with a recorded clinical dementia rating (CDR), Mini-Mental State Examination (MMSE), or postmortem clinical chart review CDR score within two years of death [22, 47]. CDR and MMSE scores were used to assign subjects into either cognitively normal or cognitively impaired groups. Individuals who had a CDR score of 0.5 or above or MMSE score below 26 were considered to be cognitively impaired while subjects with a CDR score of 0 or MMSE score 26 or above were considered cognitively normal [39]. If an individual had both MMSE score and CDR score, the most recent score was used, and if both scores were given on the same date, the CDR score was used.

Comprehensive neuropathological assessments were performed at the contributing institutions. Neuropathological criteria for PART included (1) cases that had a Braak stage of 0-IV and (2) Consortium to Establish a Registry for Alzheimer’s Disease (CERAD) neuritic plaque severity score of 0 [10, 44]. Neuropathological exclusion criteria consisted of other neurodegenerative diseases including AD, Lewy body disease, progressive supranuclear palsy (PSP), corticobasal degeneration (CBD), chronic traumatic encephalopathy (CTE), Pick disease, Guam amyotrophic lateral-sclerosis-parkinsonism-dementia, subacute sclerosing panencephalitis, globular glial tauopathy. Data pertaining to Braak stage, CERAD,

Lewy body pathology (incidental), cerebrovascular disease, infarcts (vascular brain injury), microinfarcts, and argyrophilic grains, were derived from neuropathologic studies performed at respective centers. The presence of aging-related tau astrogliopathy (ARTAG) was determined on p-tau immunohistochemical stains described below [38].

### Atrophy score

Given that no widely accepted validated system for assessing hippocampal atrophy on human brain sections exists, we devised a semiquantitative scoring system and applied it to low power images of hematoxylin & eosin-stained sections counterstained with Luxol fast blue. We defined atrophy severity as the magnitude of ventricular dilatation (hydrocephalus ex vacuo) relative to the size of the hippocampal formation. If there was no apparent ventricular dilatation or atrophy, then a score of 0 was assigned. If there was appreciable atrophy, but the dorsoventral height of the ventricle was less than the height of the thickest section of CA1, then a score of 1 (mild) was assigned. If the magnitude of ventricular dilatation exceeded the thickness of CA1, then a score of 2 (moderate) was given. If the total area of the ventricle area was greater than the area of the hippocampus proper, a score of 3 (severe) was assigned. This score was derived only in the subset of cases where the entire temporal horn of the lateral ventricle was available included in the provided section (*n*=24).

### Immunohistochemistry

Immunohistochemistry (IHC) and hematoxylin & eosin (H&E) stains were performed on FFPE sections (5 μm) that were prepared from blocks of hippocampus and frontal cortex for supplemental neuropathological analyses (see below). Sections mounted on positively charged slides were dried overnight at room temperature. IHC was performed on a Leica Bond III automated stainer, according to the manufacturer’s protocols (Leica Microsystems, Buffalo Grove, IL, USA). IHC was performed using antibodies to hyper-phosphorylated tau (p-tau, AT8, 1:1000, Fisher Scientific, Waltham, MA) and beta-amyloid (Aβ, 6E10, 1:1000, Covance, Princeton, NJ, USA). Aβ stains were confirmed to be negative to ensure that there were no neuritic or diffuse plaques present (CERAD score of 0) for all cases. For each set of slides stained, a known severe AD case was included as a batch control.

### Computer-assisted morphometric analysis

Whole slide images (WSI) were prepared from glass slides that were scanned using an Aperio CS2 (Leica Biosystems, Wetzlar Germany) digital slide scanner. Quantitative analysis of the tau burden was performed in selected regions in the hippocampi using the following methodology; WSI were neuroanatomically segmented using Aperio ImageScope software into the hippocampus proper (i.e., dentate, cornu ammonis, and subiculum) and the adjacent cortex that we termed the entorhinal region, which variably includes posterior portions of the parahippocampal gyrus with remnants of the (trans-)entorhinal region or lingual gyrus. Staining was measured in these areas separately and together using a modified version of the Aperio positive pixel count (Version 9) based on the intensities of the positive control sample in each batch to determine the area of immunoreactivity. Data were normalized using the number of positive pixel counts to the total area creating a 0-1 p-tau burden scale.

### Genetic analysis

High-throughput isolation of DNA was performed using the MagMAX DNA Multi-Sample Ultra 2.0 Kit on KingFisher Flex robotic DNA isolation system (Thermofisher, Waltham, MA). 20-40 mg of fresh frozen brain tissue were placed into a deep-well plate and treated with 480 ul of Proteinase K mix (Proteinase K, Phosphate Buffered Saline [pH 7.4], Binding Enhancer) and incubated overnight at 65°C at 800 rpm on a shaking plate. Genomic DNA was isolated and purified using magnetic particles. DNA quality control was performed using a nanodrop spectrophotometer (concentration > 50ng/ul, 260/280 ratio 1.7-2.2).

Genotyping was performed using single nucleotide polymorphism (SNP) microarrays (Infinium Global Screening Array v2.4. or the Infinium OmniExpress-24, Illumina, San Diego CA). Raw genotype files were converted to PLINK-compatible files using GenomeStudio software (Illumina, San Diego CA). *MAPT* haplotype was determined using the rs8070723 H2 tagging SNP. *APOE* genotype was provided by the collaborating center. For analyses, the *APOE* status was collapsed into a binary variable of the presence or absence of APOE ε4.

### Statistical analysis

All statistical tests were performed using the statistical software Statistical Package for the Social Sciences (SPSS) (IBM, Chicago, Il). Data was visualized using the ggplot2 package in project R or Excel (Microsoft, Redmond, Washington). Binary measurements (yes/no) were created for pathological, clinical, demographic, and genetic variables. Specifically, variables were extracted from the pathological diagnosis and binary measurements (yes/no) were created for the following variables: argyrophilic grains, Lewy body pathology (incidental), cerebrovascular disease, and infarcts (vascular brain injury). Additionally, the same process was done for clinical variables: history of psychiatric illness and education (for this study, defined as at least some college).

Descriptive statistics were used to identify differences between the cognitively normal and cognitively impaired PART groups for clinical, pathological, and genetic variables. Differences were detected using χ2 tests or exact χ2 if any cell size included < 5 participants. A t-test was performed to determine if age differed significantly between normal and cognitively groups. Next, an unadjusted binary logistic regression was performed to determine what genetic, clinical, and pathological variables were associated with being cognitively impaired within our PART cohort. Lastly, a multivariable model was created to determine what extent Braak NFT stage and the computer-assisted morphometrics were able to predict cognitive impairment in PART when adjusting for age. Statistical significance was determined if α < 0.05. Not all data was available on the subjects.

## Results

One hundred seventy-four neuropathologically confirmed amyloid-negative subjects were included in this study (Table 1, Figure 1). The overall mean age was 83.2 with a range of 52.9-105.1 years. Of these, 124 subjects (mean age 81.0, range = 52.9-102.4) had no cognitive impairment and 50 (mean age 88.3, range = 69.8-105.1) had some degree of cognitive impairment, with either mild cognitive impairment (MCI) or dementia. The majority of subjects who were cognitively impaired were 80+ years of age (Figure 2). The Braak NFT stage ranged from 0 to IV with the majority of cognitively impaired subjects having a Braak NFT score of II to IV. A higher percentage of females had cognitive impairment (62.0%) compared to those who were cognitively normal (49.2%).

**Table 1.** Patient data.

|  | Overall | Cognitive Status |  | p |
| --- | --- | --- | --- | --- |
|  |  | Normal | Impaired* |  |
| *Demographics* |  |  |  |  |
| Average age at testing (range) | 83.2 (52.9-105.1) | 81.0 (52.9-102.4) | 88.3 (69.8-105.1) | **<0.0001** |
| Total (Male / Female) | 174 (82 / 92) | 124 (63 / 61) | 50 (19 / 31) | 0.126*** |
| Age at last visit (%) |  |  |  |  |
| <60 | 7 (4.0) | 7 (5.6) | 0 (0.0) |  |
| 60-69 | 15 (8.6) | 14 (11.3) | 1 (1.7) |  |
| 70-79 | 33 (19.0) | 30 (24.2) | 3 (5.2) |  |
| 80-89 | 76 (43.7) | 45 (36.3) | 31 (53.4) |  |
| 90+ | 51 (29.3) | 28 (22.6) | 23 (39.7) |  |
| Education, at least some college (%) | 32 (18.4) | 15 (78.9) | 17 (77.3) | 0.89 |
| History of psychiatric illness (%) | 45 (25.9) | 29 (31.9) | 17 (45.9) | 0.13 |
| *Neuropathological data* |  |  |  |  |
| Argyrophilic grains | 32 (18.4) | 12 (9.7) | 10 (20.0) | 0.06 |
| Lewy body pathology (incidental) | 16 (9.2) | 11 (8.9) | 5 (10.0) | 0.82 |
| Cerebrovascular disease** | 27 (15.5) | 6 (4.8) | 21 (42.0) | **<0.0001** |
| Infarcts (vascular brain injury) | 37 (21.3) | 24 (19.4) | 13 (26.0) | 0.33 |
| Hippocampus ARTAG positive (%) | 43 (24.7) | 25 (21.6) | 18 (38.3) | **0.03** |
| *Genetic Data* |  |  |  |  |
| Presence of ≥1 APOE ε4 allele | 22 (12.6) | 16 (12.9) | 6 (11.3) | 0.77 |
| Presence of ≥1 APOE ε2 allele | 46 (26.4) | 27 (21.8) | 19 (35.8) | 0.06 |
| Presence of ≥1 MAPT H2 | 59 (33.9) | 42 (36.2) | 17 (36.2) | 1 |

\* Mild cognitive impairment or dementia, \*\* excluding cerebral amyloid angiopathy, \*\*\*Male sex, significant values in bold (Chi squared test)

**Figure 1.**
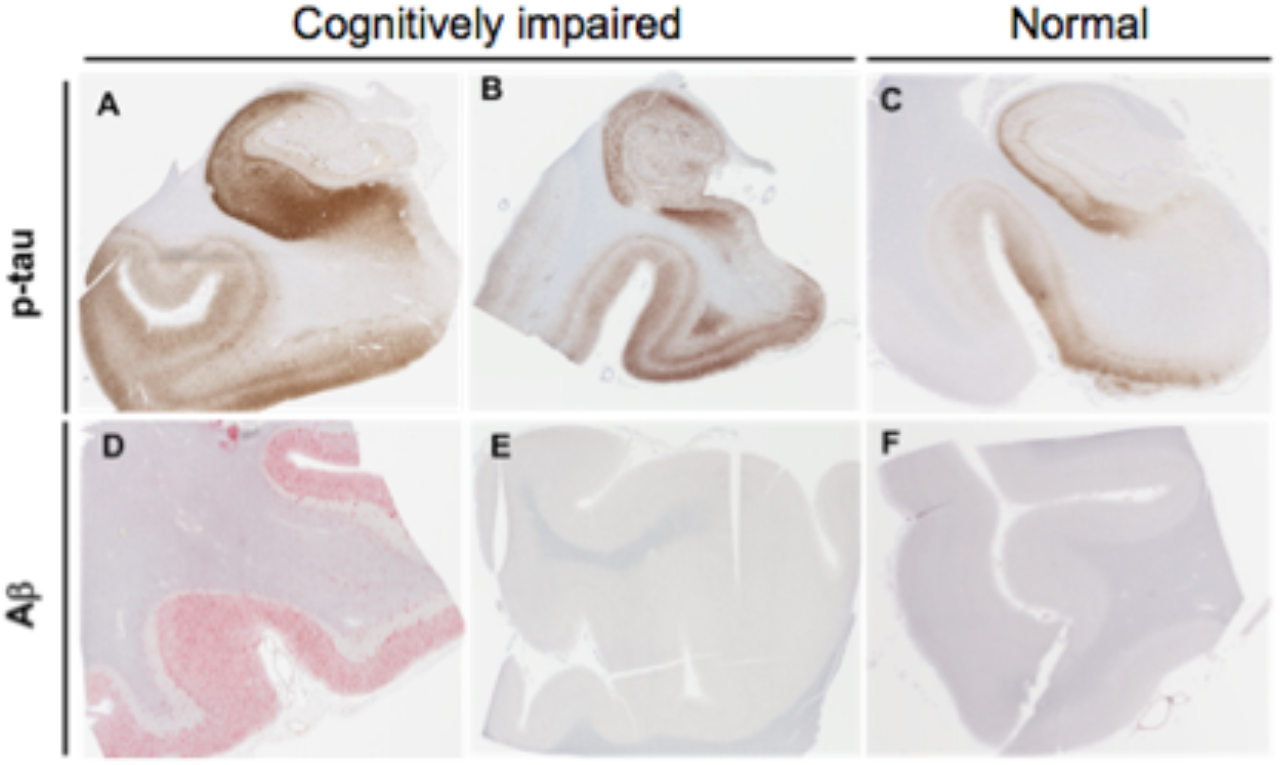
Comparison of amyloid and tau pathology in primary age-related tauopathy (PART) versus Alzheimer disease (AD). **(A) Immunohistochemical staining using antisera to hyperphosphorylated tau in** an AD brain shows marked hyperphosphorylated tau (p-tau)-containing neurofibrillary tangles (NFT) in the hippocampus which extends past the collateral sulcus into the parahippocampal gyrus and other neocortical regions. **(B, C)** Subjects with mild to severe PART have elevated p-tau levels in the hippocampus predominantly restricted to the medial temporal lobe. (**D, E, F**) Subjects with AD neuropathologic change have abundant Aβ-containing plaques in neocortical structures, whereas those with PART have sparse or none. These neuropathologic changes in AD and PART are seen in association with varying degree of cognitive impairment ranging from cognitively normal to demented.

**Figure 2.**
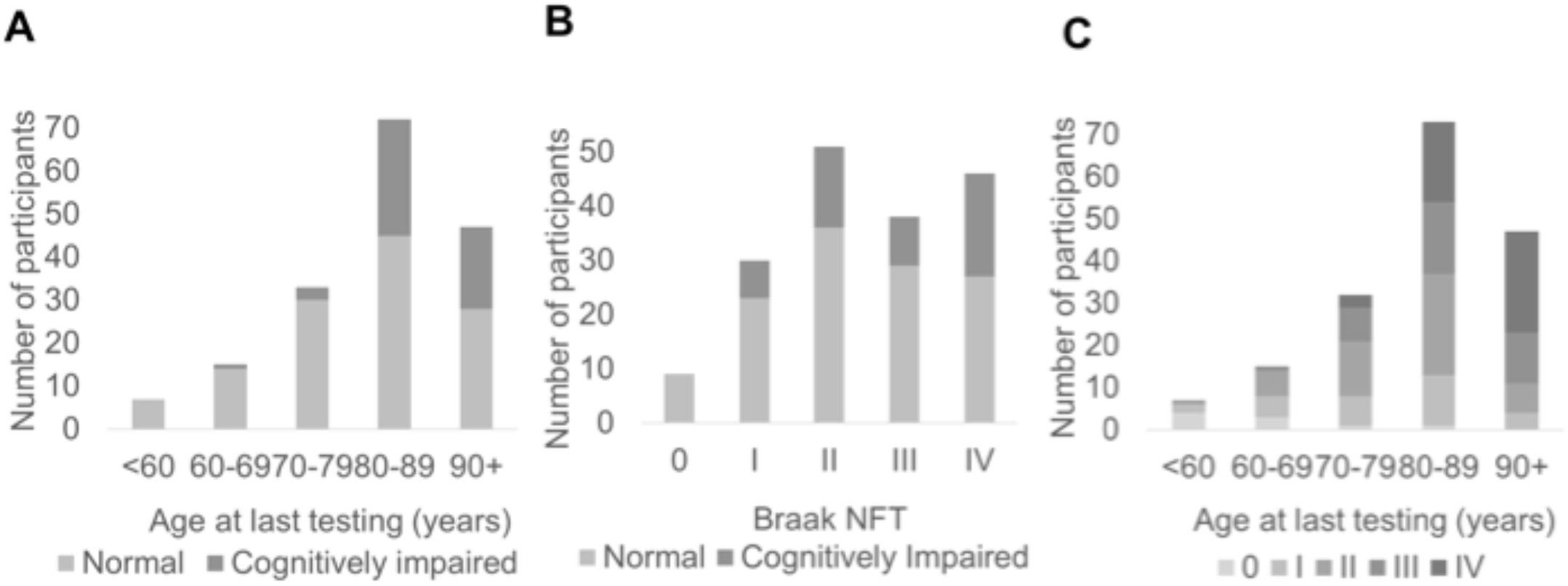
Distribution of age, Braak neurofibrillary tangle (NFT) stage and cognitive status. (**A**) The number of normal and cognitively impaired subjects across the age spectrum. (**B**) The number of cognitively normal and impaired subjects by Braak stage. (**C**) The number of subjects across the aging spectrum by Braak stage.

We observed several differences among subjects with cognitive impairment compared to those who were cognitively normal. First, cognitively impaired PART subjects were more likely to be older (age of testing 81.0 vs. 88.3, *p* < 0.0001), have cerebrovascular disease (42.0% vs. 4.8%, *p* < 0.0001) and have hippocampal age-related tau astrogliopathy (ARTAG; 38.3% vs. 21.6%, *p* < 0.05) compared to cognitively normal subjects (Table 1). However, education, history of psychiatric illness, argyrophilic grains, incidental Lewy body pathology, infarcts, presence of an *APOE* ε2 allele, presence of *APOE* ε4 allele, and *MAPT* haplotype status did not significantly affect cognitive status (*p* > 0.05 for all conditions).

In our main unadjusted analysis, we assessed the extent to which a series of clinical, neuropathological, and genetic variables predicted cognitive impairment in our PART cohort (Table 2). We found that age and cerebrovascular disease were the strongest predictors of cognitive impairment (*p* < 0.0001 for both cases). ARTAG and hippocampal atrophy were also significant predictors, but to a lesser extent (*p* < 0.05 for both cases). There were more reported men and subjects with a history of psychiatric illness, argyrophilic grains, incidental Lewy body pathology, infarcts, and microinfarcts in the cognitively impaired PART group, however none of these predictors was significantly different (*p* > 0.05 for all conditions). *APOE* ε4 (at least 1 ε4 allele) was reported more in the cognitively normal PART group but did not reach significance. Braak NFT stage significantly predicted cognitive impairment (*p* < 0.05).

**Table 2.** Unadjusted odds of being cognitively impaired.

|  | OR | 95% CI | p value |
| --- | --- | --- | --- |
| *Characteristic* |  |  |  |
| Age, at testing | 1.08 | 1.04-1.13 | **<0.0001** |
| Education, y | 0.87 | 0.67-1.12 | 0.28 |
| Sex | 1.69 | 0.86-3.30 | 0.13 |
| APOE (at least 1 ε4 allele) | 0.988 | 0.36-2.70 | 0.98 |
| History of psychiatric diagnosis | 1.82 | 0.83-3.98 | 0.14 |
| Aging-related tau astroglipathy (ARTAG) | 2.26 | 1.08-4.72 | **0.03** |
| Argyrophilic grains | 2.33 | 0.94-5.82 | 0.07 |
| Lewy body pathology (incidental) | 1.14 | 0.38-3.47 | 0.82 |
| Cerebrovascular disease* | 14.24 | 5.27-38.48 | **<0.0001** |
| Infarcts (vascular brain injury) | 1.46 | 0.68-3.17 | 0.33 |
| Microinfarcts | 1.05 | 0.43-2.59 | 0.91 |
| Hippocampal atrophy | 5.32 | 1.04-27.09 | **0.04** |
| Braak NFT stage | 1.37 | 1.03-1.83 | **0.03** |
| *Computer-assisted p-tau (AT8) burden (positive pixel counts)* |  |  |  |
| Entorhinal region | 1.90 | 1.31-2.75 | **0.001** |
| Hippocampus proper | 2.17 | 1.48-3.20 | **<0.0001** |
| Entorhinal region & Hippocampus proper | 2.12 | 1.44-3.11 | **<0.0001** |

\* Excluding cerebral amyloid angiopathy, significant values in bold (logistic regression)

Additionally, the computer-assisted morphometrics in the entorhinal region, hippocampus proper, and the combined region were significantly associated with cognitive impairment (*p* = 0.0001, Figure 3A-C, Table 3). Lastly, when the Braak NFT stage was correlated with computer-assisted morphometrics in the combined region (*p* < 0.001), there was a high degree of variability between the Braak NFT stage and the computer-assisted combined region morphometrics (Figure 3D).

**Figure 3.**
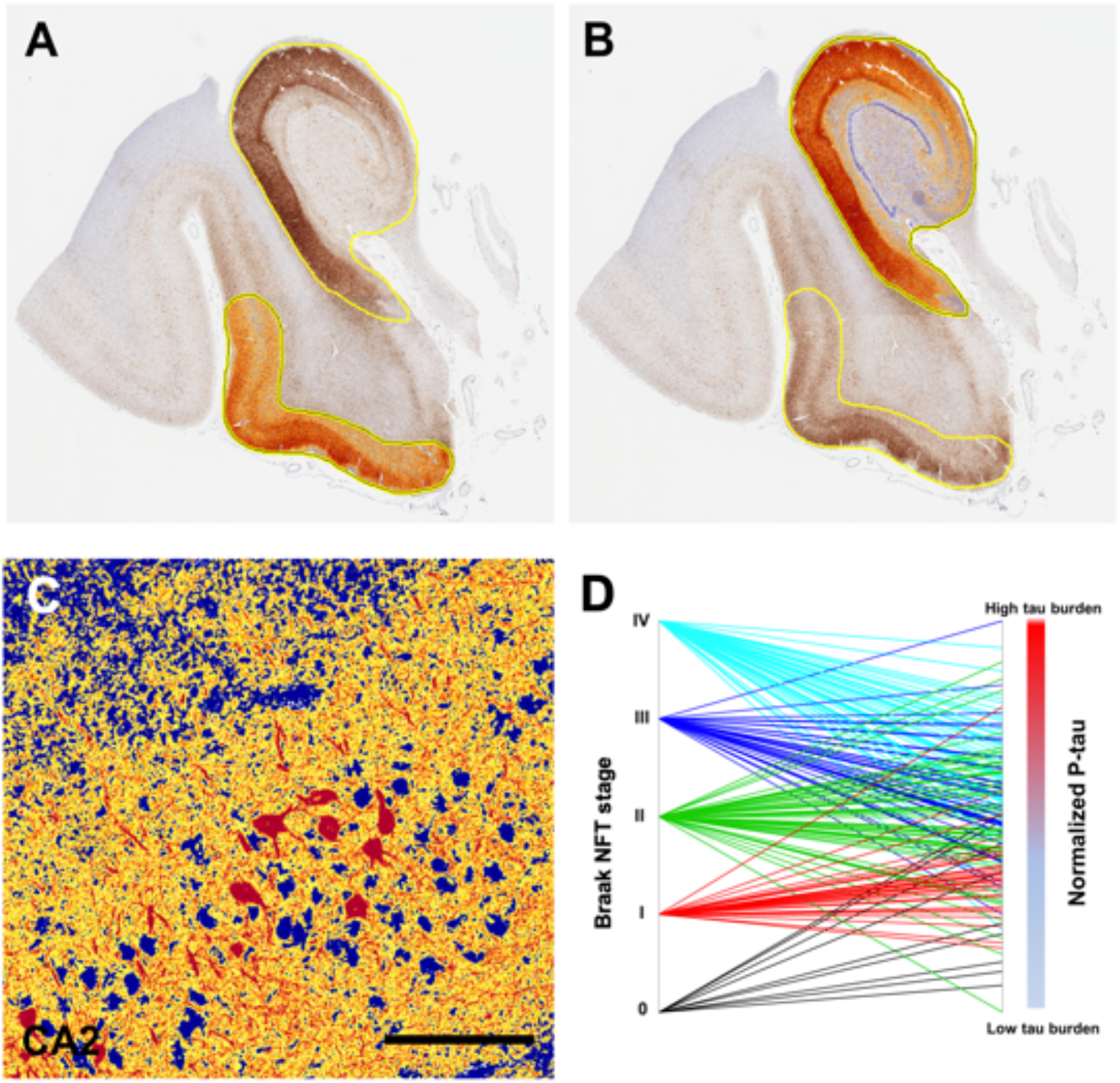
Computer-assisted morphometrics to assess pathological tau burden. (**A, B**) Quantitative assessment of hyperphosphorylated tau (p-tau) burden was performed on whole slide images of the hippocampus stained for p-tau (AT8) using immunohistochemistry. Positive pixel counts were determined in two regions (hippocampus proper and entorhinal region). Results were normalized to the total area assessed. A third summary score of the total p-tau burden of the medial temporal lobe was calculated by summing positive pixels in both. (**C**) High power image shows high intensity in red, medium intensity in yellow and negative staining in blue. (**D**) Parallel plot showing the relationship between Braak stage and the computer morphometric quantification of p-tau using the normalized medial temporal lobe (hippocampus and entorhinal region). Scale bar = 150 μm.

**Table 3.** Odds of being cognitively impaired at death, adjusted.

|  | **OR** | **95% CI** | ***p* value** |
| --- | --- | --- | --- |
| Braak NFT stage | 1.01 | 0.72-1.41 | 0.98 |
| P-tau burden (computer-assisted AT8 IHC positive pixels) |  |  |  |
| Entorhinal region | 1.46 | 0.97-2.20 | 0.07 |
| Hippocampus | 1.66 | 1.07-2.57 | **0.02** |
| Entorhinal region & hippocampus | 1.62 | 1.06-2.49 | **0.03** |
| Significant values in bold (logistic regression) |  |  |  |

Finally, using a multivariable model, we assessed whether any measurements for p-tau predicted cognitive impairment when controlling for age. In this adjusted analysis, we found that computer-assisted morphometrics used to capture p-tau burden in the hippocampus proper and combined region were significantly associated with cognitive impairment in PART (*p* < 0.05 for both cases). However, the computer-assisted morphometrics in the entorhinal region were not associated with cognitive impairment yet there was a trend toward statistical significance (*p* = 0.068). The Braak NFT stage was not able to predict cognitive impairment when controlling for age (*p* = 0.73, Table 3, Figure 4).

**Figure 4.**
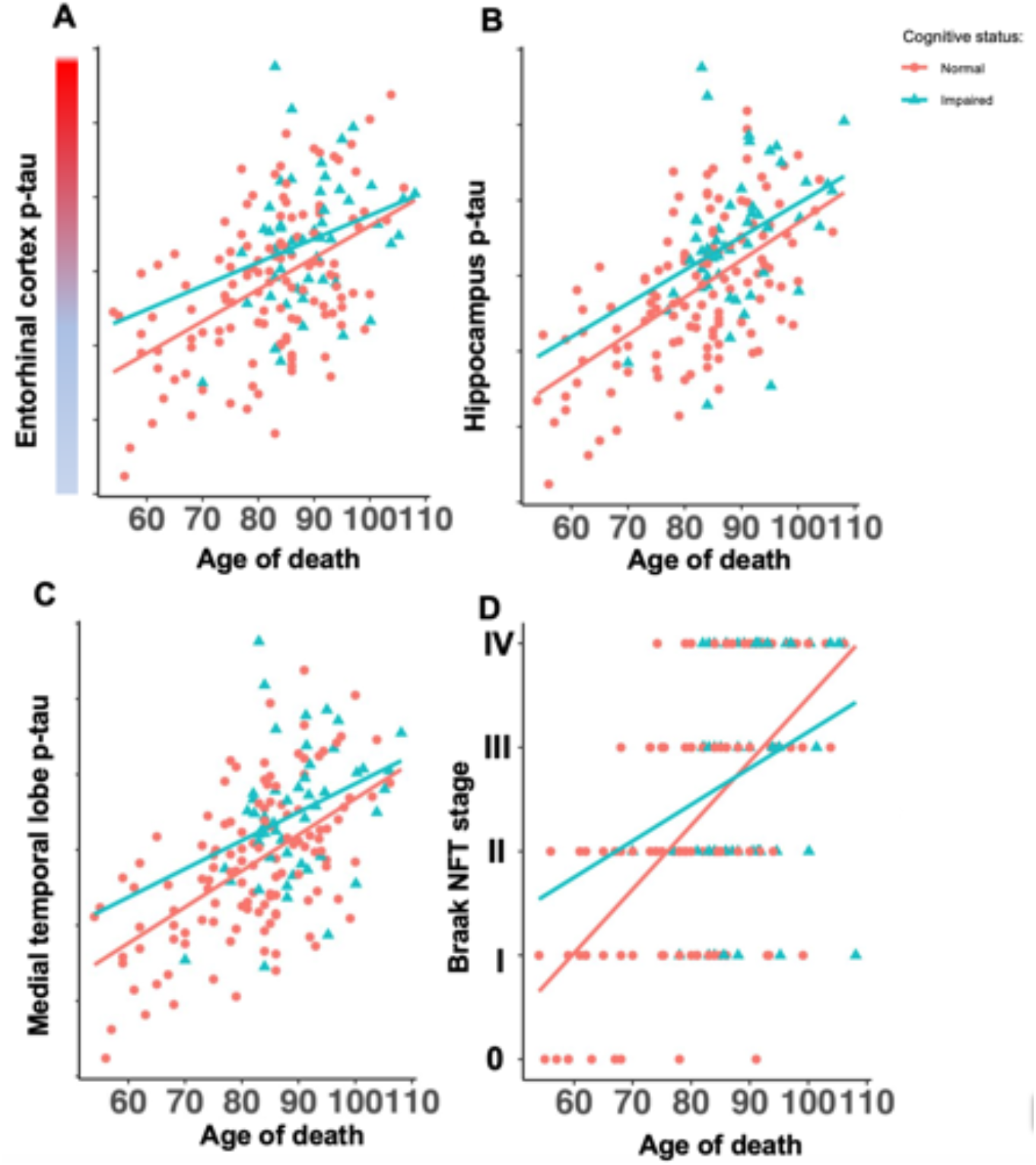
Pathological tau burden in normal and cognitively impaired subjects across the aging spectrum. (**A-C**) Generalized linear models of age versus tau burden show significant differences between cognitively normal and cognitively impaired subjects in the hippocampus proper (*p* = 0.047), and combined entorhinal region and hippocampus regions (*p* < 0.048), but not in the entorhinal region alone (*p* = 0.07). (**D**) Generalized linear model of age vs Braak NFT staging did not show significant differences between cognitively normal and cognitively impaired subjects (*p* = 0.73).

## Discussion

Since the neuropathological criteria for PART were proposed, the terminology has been widely adopted, but controversy persists, especially around its relationship to Alzheimer disease (AD). Delineating the histological/cellular features that are associated with cognitive impairment in PART is critical for advancing our understanding of the pathology and determining the extent to which it overlaps with AD. The fact that subjects with PART, as with AD neuropathologic change, can range in their cognitive status from normal to demented, raises the question as to whether cognitive reserve/resilience plays a role or alternatively whether we are not adequately capturing the relevant features, such as common comorbidities or other factors. This study, by using a large autopsy cohort with multivariate analyses, directly addresses these critical questions. The goal was to leverage our collection of post-mortem PART brains to characterize the clinical, pathological, and genetic features that are associated with cognitive impairment in PART. Additionally, we sought to compare Braak stage with pathology burden measures derived from p-tau immunohistochemistry that quantifies severity independently of neuroanatomical vulnerability. To overcome intra-center variability in tau pathology measures, we reassessed each case histologically to maximize accuracy.

We found that all of our PART definite cases had p-tau restricted mainly to the MTL (Braak NFT stage <IV), which is consistent with and supports other previous studies investigating PART [4, 17, 34]. Cases ranged in cognitive impairment with the majority of subjects being cognitively normal, and consistent with prior data, the PART subjects tended to be older than individuals with AD [17, 59]. The results of our study confirm those of previous autopsy studies showing that cerebrovascular disease predicts cognitive impairment in PART [6, 49]. Interestingly, we did not see a strong correlation between cognitive impairment and microinfarcts, while others have shown a correlation with cognition in the oldest old [14]. We did however, find novel, unreported associations of increased age, hippocampal atrophy, and ARTAG with cognitive impairment in our PART definite cohort. Similar to what has been reported by those utilizing the NACC database, our results verify those with a higher Braak NFT stage are associated with more rapid cognitive decline [33].

While these associations have yet to be reported in PART, there are numerous studies showing that age, atrophy, and ARTAG may be associated with cognitive impairment [9, 23, 32, 50, 53]. Surprisingly, we did not see increased odds of the Braak NFT stage being associated with cognitive impairment when controlling for age as has been reported in other studies [6]. However, we did find that using computer-assisted morphometrics to assess p-tau burden in the entorhinal region, hippocampus, and combined region was able to significantly predict cognitive impairment, similar to other studies [1, 13]. While Braak NFT staging is the most widely employed approach for assessing p-tau, it is limited in that it primarily focuses on regionality and not disease burden [25]. Other studies have employed both manual and computer assisted quantitative approaches that may capture aspects of pathological features with more power [24, 26, 56]. However, a majority of these approaches focuses on AD which may not be relevant in the context of PART, where p-tau pathology does not progress in the same hierarchical manner proposed by Braak in AD [10, 16]. Hence this study highlights several new methodologies to assess p-tau burden, which our results suggest to be a more accurate predicator of clinical symptomology in those with PART.

In addition to assessing p-tau burden, we also examined the effect of *APOE* status in PART as a predictor for cognitive impairment. *APOE* ε4 has been strongly suggested as an important predictor of cognitive decline in AD while *APOE* ε2 has been shown to be protective [15, 18, 54, 58]. However, many of these studies have been performed in AD cohorts, and in aging cohorts there has been evidence suggesting the ε4 allele is not a risk factor for cognitive impairment [57]. Our data agree with that reported by Small *et al*. as we did not see an association with *APOE* ε4 and cognitive impairment, which might be explained by the fact that we studied a pathologically confirmed amyloid-negative cohort. Recent work has suggested that *APOE* may exacerbate tau pathology independently of amyloid deposition [55]. Here, we failed to detect an association of cognitive impairment in PART with the *MAPT* H1 haplotype; future larger studies with more statistical power are required to delineate the genetic architecture of PART.

This study had notable limitations. There was a relatively small number of subjects in the cognitively impaired PART group (*n*=50), which may weaken our power to predict cognitive impairment.

Additionally, because a majority of our subjects were not from longitudinally studied prospective cohorts, we were unable to obtain certain lifestyle variables, such as actual years of education and concussion history, which could potentially significantly affect our model. However, given that diagnosing PART pre-mortem is currently challenging, it would be impractical to create such a prospective cohort. We would also like to highlight the association we observed with ARTAG and cognitive status might be only due to collinearity between p-tau severity and ARTAG, with p-tau probably the driving pathology and the ARTAG association being significant because of its potential dependence on p-tau. Lastly, our study was limited to pathology of the medial temporal lobe and frontal cortex. A more exhaustive study would have incorporated a greater number of brain regions to more extensively address other potential tau-related pathologies.

In summary, our findings are consistent with the hypothesis that PART is an amyloid-independent tauopathy, primarily affecting the medial temporal lobe, which can present with cognitive impairment. Several demographic and neuropathological variables including age, ARTAG, cerebrovascular disease, hippocampal atrophy, Braak NFT stage, and p-tau computer assessments were significantly associated with cognitive impairment in our PART cohort. The Braak NFT stage was not a significant predictor of cognitive impairment when controlling for age, while the computer-assistant morphometrics were. These data strongly suggest that neuroanatomical staging used in AD may not be as relevant to PART given the pathology minimally spreads beyond the medial temporal lobe. Novel techniques to measure p-tau burden can further our understanding of PART pathology and associated clinical and genetic features.

## Acknowledgements

This work was supported by the National Institutes of Health [R01 AG054008, R01 NS095252, R01 AG060961, and R01 NS086736 to J.F.C, F32 AG056098 and P30 AG066514 to K.F., R01 AG062348 to J.F.C., A.M., and D.D., P30 AG010124, P01 AG017586 and U19 AG062418 to J.Q.T, R01 AG066152 to C.T.M, P50 AG005133 to J.K., P50 AG005138, P30 AG066514, and 75N95019C00049 to V.H., U24 NS072026 and P30 AG019610 to T.B., P30 AG013854 to E.B., P30 NS055077 and P50 AG025688 to M.G., P30 AG08017 to R.W. and U54 NS115266 to A.M.], the Alzheimer’s Association [NIRG-15-363188 to J.F.C.], the Tau Consortium, Genentech/Roche, David & Elsie Werber, Alexander Saint-Amand Fellowship, J.M.R. Barker Foundation, The McCune Foundation, and the Winspear Family Center for Research on the Neuropathology of Alzheimer Disease, The Arizona Department of Health Services, and the Michael J. Fox Foundation for Parkinson’s Research. G.G.K. is supported by the Rossy Foundation and by the Safra Foundation. The authors would also like to acknowledge Ping Shang, HT(ASCP) QIHC and Jeff Harris, HTL(ASCP) for histologic and immunohistochemical preparations, and Chan Foong, M.S., for preparation of whole slide image.

## Notes

The authors declare no conflicts of interest

### Competing Interest Statement

The authors have declared no competing interest.

